# Rapid alignment-free phylogenetic identification of metagenomic sequences

**DOI:** 10.1101/328740

**Authors:** Benjamin Linard, Krister Swenson, Fabio Pardi

## Abstract

**Motivation:** Taxonomic classification is at the core of environmental DNA analysis. When a phylogenetic tree can be built as a prior hypothesis to such classification, phylogenetic placement (PP) provides the most informative type of classification because each query sequence is assigned to its putative origin in the tree. This is useful whenever precision is sought (e.g. in diagnostics). However,likelihood-based PP algorithms struggle to scale with the ever-increasing throughput of DNA sequencing.

**Results:** We have developed RAPPAS (Rapid Alignment-free Phylogenetic Placement via Ancestral Sequences) which uses an alignment-free approach, removing the hurdle of query sequence alignment as a preliminary step to PP. Our approach relies on the precomputation of a database of k-mers that may be present with non-negligible probability in relatives of the reference sequences. The placement is performed by inspecting the stored phylogenetic origins of the k-mers in the query, and their probabilities. The database can be reused for the analysis of several different metagenomes. Experiments show that the first implementation of RAPPAS is already faster than competing likelihood-based PP algorithms, while keeping similar accuracy for short reads. RAPPAS scales PP for the era of routine metagenomic diagnostics.

**Availability:** Program and sources freely available for download at gite.lirmm.fr/linard/RAPPAS.

## 1. Introduction

Inexpensive high-throughput sequencing has become a standard approach to inspect the biological content of a sample. Well-known examples range from health care, where sequencing-based diagnostics are being implemented in many hospitals and in reaction to disease outbreaks (Gilchrist et al., 2015; Gardy and Loman, 2018), to environmental and agricultural applications, where the diversity of the microorganisms in soil, water, plants, or other samples can be used to monitor an ecosystem (Deiner et al., 2017; Porter and Hajibabaei, 2018). Furthermore, with the advent of sequencing miniaturization, the study of environmental DNA will soon be pushed to many layers of society (Gilbert, 2017; Zaaijer et al., 2016).

Metagenomics is the study of the genetic material recovered from this vast variety of samples. Contrary to genomic approaches studying a single genome, metagenomes are generally built from bulk DNA extractions representing complex biological communities that include known and yet unknown species (*e.g.* protists, bacteria, viruses). Because of the unexplored and diverse origin of the DNA recovered, and because of the sheer volume of sequence data, the bioinformatic analysis of metagenomes is very challenging. In particular, identifying the species at the origin of metagenomic sequences remains a bottleneck for analyses based on environmental DNA.

Several classes of approaches are possible for taxonomic classification of metagenomic sequences, the choice of which will generally depend on the availability of prior knowledge (*e.g.* a reference database of known sequences and their relationships) and a trade-off between speed and precision. For instance, without prior knowledge, the exploration of relatively unknown clades can rely on unsupervised clustering (Mahé et al., 2014; Ondov et al., 2016; Sedlar et al., 2017). The precision of this approach is limited, and clusters are generally assigned to taxonomic levels via posterior analyses involving local alignment searches against reference databases. An alternative is to contextualise local alignments with taxonomic metadata. For instance, MEGAN (Huson et al., 2007, 2016) assigns a query metagenomic sequence (QS for simplicity) to the last common ancestor (LCA) of all sequences to which it could be aligned with a statistically-significant similarity score (*e.g.* using BLAST or other similarity search tools). A faster alternative is to build a database of taxonomically-informed k-mers which are extracted from a set of reference sequences. Classification algorithms then assign a QS to the taxonomic group sharing the highest number of the k-mers (Ames et al., 2013; Wood and Salzberg, 2014; Müller et al., 2017; Liu et al., 2018). All of these approaches use simple sequence similarity metrics and no probabilistic modeling of the evolutionary phenomena linking QSs to reference sequences. The advantage is that, particularly in the case of k-mer-based approaches, these methods facilitate the analysis of millions of QSs in a timescale compatible with large-scale monitoring and everyday diagnostics.

Nevertheless, in many cases a precise identification of the evolutionary origin of the QSs is desired. This is particularly true for medical diagnostics where adapted treatments will be administered only after precise identification of a pathogen (Butel, 2014; Trémeaux et al., 2016). In the case of viruses (poorly known and characterized by rapid evolution), advanced evolutionary modelling is necessary to perform detailed classifications. A possible answer in this direction may be the use of phylogenetic inference methods. However, the standard tools for phylogenetic inference are not applicable to metagenomic datasets, simply because the sheer number of sequences cannot be handled by current implementations (Izquierdo-Carrasco et al., 2011).

A solution to this bottleneck is *phylogenetic placement* (PP) (Matsen et al., 2010; Berger et al., 2011). The central idea is to ignore the evolutionary relationships between the QSs and to focus only on elucidating the relationships between a QS and the known reference sequences. A number of reference genomes (or genetic markers) are aligned and a “reference” phylogenetic tree is inferred from the resulting alignment. This reference tree is the backbone onto which the QSs matching the reference genomes will be “placed”, *i.e.* assigned to specific edges. When a QS is placed on an edge, this is interpreted as the evolutionary point where it diverged from the phylogeny. Note that when the placement edge of a QS is not incident to a leaf of the tree, the interpretation is that it does not belong to any reference genome, but to a yet unknown/unsampled genome for which the divergence point has been identified. This is different from LCA-based approaches where a QS assigned to an internal node usually means that it cannot be confidently assigned to any leaf of the corresponding subtree. Currently, only two PP software implementations are available: PPlacer (Matsen et al., 2010; McCoy and Matsen, 2013) and EPA (until recently a component of RAxML) (Berger et al., 2011). They are both *likelihood-based*, *i.e.* they try to place each QS in the position that maximises the likelihood of the tree that results from the addition of the QS at that position. Alternative algorithms were also described, but not yet publicly released (Brown and Truszkowski, 2013; Filipski et al., 2015). These methods have some limitations: (i) likelihood-based methods require an alignment of each QS to the reference genomes prior to the placement itself, a complex step that may dominate the computation time for long references, and (ii) despite being much faster than a full phylogenetic reconstruction, they struggle to scale with increasing sequencing throughputs. Recently, the development of EPA-ng (Barbera et al., 2018) provided an answer to the second point via the choice of advanced optimization and efficient parallel computing. But no fundamental algorithmic changes were introduced, and the method still relies on a preliminary alignment.

To avoid these bottlenecks, we have developed the phylogenetic placement program RAPPAS (Rapid Alignment-free Phylogenetic Placement via Ancestral Sequences). The novelty of our approach rests on two key ideas: (i) a database of “phylogenetically-informed” k-mers is built from the reference tree and reference alignment, (ii) QSs are then placed on the tree by matching their k-mers against the phylogenetic k-mer database. For a specific group of organisms, the maintenance of the RAPPAS database can be seen as a periodic “build-and-update” cycle, while the phylogenetic placement of new metagenomes becomes a scalable, alignment-free process compatible with day-to-day routine diagnostics. This algorithmic evolution bypasses the alignment of QSs to the reference required by existing algorithms. For short reads RAPPAS produces placements that are as accurate as previous likelihood-based methods. Despite the potential for improvement, our experiments show that RAPPAS is already significantly faster on real datasets than the most optimized versions of competing tools.

## 2. Methods

### 2.1 Analyzing metagenomes with RAPPAS

In order to perform phylogenetic placement, one must define a reference dataset (alignment and tree) onto which future QSs will be aligned and placed. Typically, in a context of metagenomic analysis and large-scale taxonomic classification, such references will be based on well-established genetic markers (*e.g. 16S rRNA, rbcl, etc.)* or short genomes (*e.g.* organelles, viruses, *etc.*).

One of the main differences between our approach and the existing PP methods is in the workflows that they entail. Existing PP algorithms require the alignment of every QS to the reference alignment prior to the placement itself, a complex step that relies on techniques such as profile Hidden Markov Models that may dominate computation time. For each sample analyzed, the alignment phase is then followed by the placement phase, which relies on phylogenetic inference (Figure 1-A). RAPPAS uses a more efficient approach where all computationally-heavy phylogenetic analyses are performed as a preprocessing phase, which is only executed once before the analysis of any metagenomic sample. This phase builds a data structure (named pkDB; see next section) which will be the only information needed to place QSs. Then many samples can be routinely placed on the tree without the need for any alignment. The same pkDB can be reused for several query datasets (Figure 1-B).

**Figure 1:**
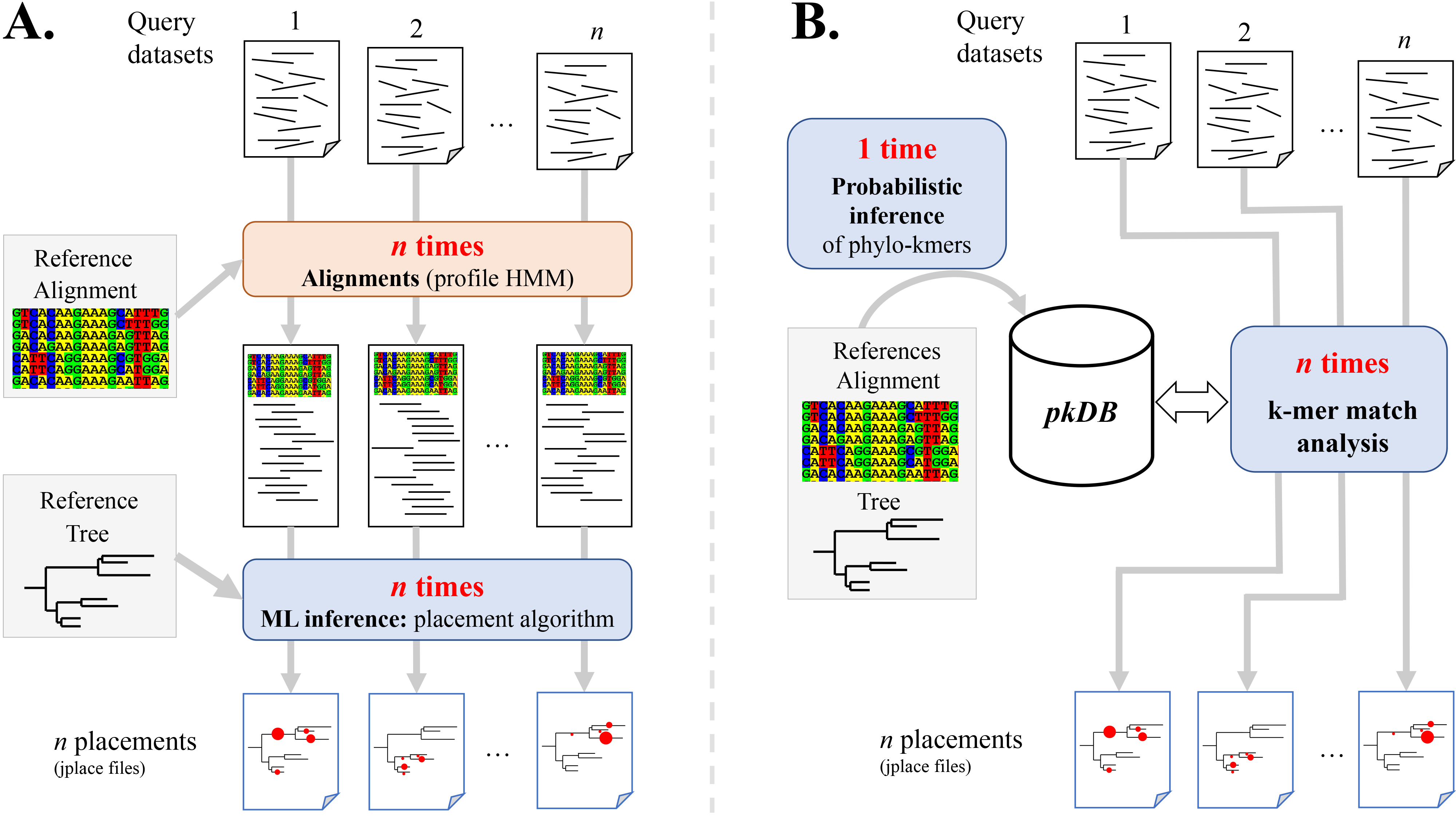
Comparison between existing PP software and RAPPAS. The pipelines from query sequence datasets to placement results (jplace files) are depicted. **A.** Likelihood-based software requires, for each new query dataset, the step of aligning the query sequences to the reference alignment via an external tool (red box). The resulting extended alignment is the input for the phylogenetic placement itself (blue box). **B.** RAPPAS builds a database of k-mers (the pkDB) once for a given reference tree and alignment. Many query datasets can then be placed without alignment, matching their k-mer content to this database. Each operation is run with a seperate call to RAPPAS (blue boxes).

An advantage of this approach is that many different pkDBs can be computed for different reference datasets and copied to less powerful devices for routine analyses. To follow the standards defined by previous PP methods, the placements are reported in a JSON file following the “jplace” specification (Matsen et al., 2012) (see Suppl. File S1 for more details). The results of RAPPAS can then be exploited in all jplace-compatible software, such as iTOL (visualization), Guppy (diversity analyses) or BoSSA (visualisation and diversity analysis) (Letunic and Bork, 2016; Matsen and Evans, 2013; Lefeuvre, 2018).

### 2.2 Construction of the phylo-kmers DB (pkDB)

Before performing placements, a database of phylogenetically informative k-mers must be built; these *phylo-kmers* are those that we would expect to see in a QS with non-negligible probability. The input for this phase is a reference alignment (denoted *refA*) and a reference tree (denoted *refT*) reconstructed from refA. The refT is assumed to be rooted and with lengths associated to its edges. This process is divided in three steps (see Figure 2): A) the injection of “ghost nodes” on every edge of refT, B) a process of ancestral state reconstruction based on this extended tree, and C) the generation of the phylo-kmers and their storage in a data structure.

**Figure 2:**
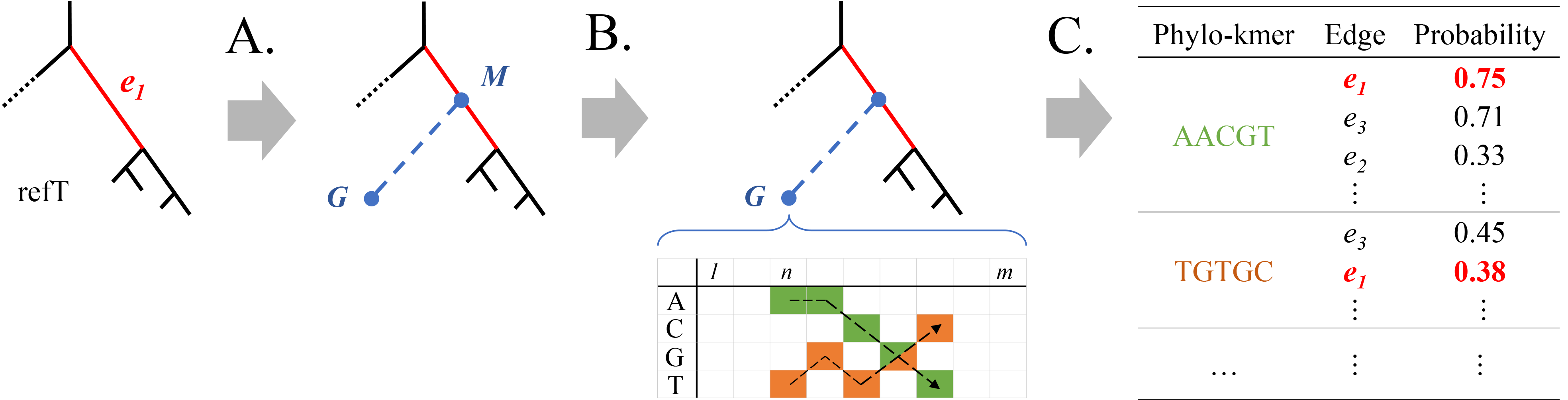
Construction of the phylo-kmers DB. The construction of phylo-kmers for an edge is depicted (see the *Methods* section for details.) **A.** Each edge *e*_1_ is bisected by a ghost node *M*, to which a terminal edge leading to another ghost node *G* is attached. **B.** For each ghost node, a probability table for the sequence states expected at each site is generated using ancestral state reconstruction. **C.** The most probable k-mers are computed from each table and then stored in the pkDB, together with the corresponding edge in the reference tree and the corresponding probability.

The first step enables us to simulate evolution that has occurred and resulted in sequences not present in the reference alignment. For each edge of refT, a new edge branching off at its midpoint *M* is created (Figure 2, step A). The length of the new edge is set to the average length over all paths from *M* to its descendant leaves. The new leaves of these new edges (e.g. *G* in Figure 2), and the midpoints *M* are referred to as *ghost nodes*. They represent unsampled sequences related to those in the reference alignment, whose most probable k-mers are the phylo-kmers. Newly sampled QSs will tend to contain phylo-kmers that are informative of their correct phylogenetic placement.

The second step computes the probabilities necessary to the construction of the phylo-kmers. It is based on standard phylogenetic methods for ancestral sequence reconstruction (Figure 2, step B). Given refA, an alignment consisting of *m* sites over 5 possible states (4 for DNA, 20 for amino acids), for each individual ghost node *G* we compute an *s* × *m* table (*p*_σ,*j*_) of probabilities, where *p*_*σ,j*_ represents the *marginal* probability of state σ at site *j* (Yang et al., 1995). From this table, we compute the probabilities of any given k-mer at any ghost node *G* (Figure 2, step C). Let *k* be a k-mer size and let *n* ∈ {1,2,…,*m* − (*k* − 1)} be a particular site of refA. Assuming independence among sites (see, *e.g*., Felsenstein, 2004), the probability *P* of k-mer σ_1_σ_2_…σ_*k*_ in *G* at sites {*n*, *n* + 1,…, *n* + *k* − 1} is given by:

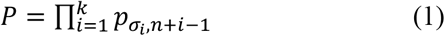

Each k-mer with a probability *P* larger than a (parameterized) threshold (by default 1/*S*^*k*^) is stored together with (i) the edge corresponding to the ghost node of origin, and (ii) the probability *P* associated to this k-mer at the ghost node of origin (Figure 2, step C).

In the end, all the k-mers reconstructed above are stored in a data structure that allows fast retrieval of all relevant information associated to a k-mer, namely the list of edges for which it has high-enough probability, and the probabilities associated to each of these edges. This structure will be referred to as the *phylo-kmers database* (*pkDB*). We note that when the same k-mer is generated at different positions *n* of refA, or at different ghost nodes for the same edge, only the highest probability is retained in the pkDB.

### 2.3 Placement algorithm

RAPPAS places each QS on the sole basis of the matches that can be found between the k-mers in the QS and the phylo-kmers stored in the pkDB. An algorithm akin to a weighted vote is employed: each k-mer in a QS casts multiple votes – one for each edge associated to that k-mer in the pkDB – and each vote is weighted proportionally to the (logarithm of the) k-mer probability at the voted edge. The edge or edges receiving the largest total weighted vote are those considered as the best placements for a given QS. A detailed description and complexity analysis for this algorithm can be found in Suppl. File S1. Unlike for alignment-dependent algorithms, the running time of RAPPAS does not depend on the length of the reference sequences, and rather than scaling linearly in the size of the reference tree, it only depends on the number of edges that are actually associated to some k-mer in the QS.

### 2.4 Accuracy experiments

The accuracy of the placements produced by RAPPAS and by competing likelihood-based PP software were evaluated following established simulation protocols (Matsen et al., 2010; Berger and Stamatakis, 2011). Briefly, these protocols rely on independent pruning experiments, in which sequences are removed from refA and refT, and then used to simulate QSs (we use the word *read* interchangeably, in the context of simulations), which are finally placed back to the pruned tree. The QSs are expected to be placed on the edge from which they were pruned. If the observed placement disagrees with the expected one, a topological metric quantifies the distance between them by counting the number of nodes separating the two edges. To make the placement procedure more challenging, we not only prune randomly chosen reference taxa, but whole subtrees chosen at random. We consider this approach realistic in the light of current metagenomic challenges, because it is indeed common that large sections of the tree of life are absent from the reference phylogenies. We also decided to simulate not just one QS per pruned sequence, but as many non-overlapping QSs as permitted by the length of the pruned sequence (*e.g.* when generating 150 bp reads, a pruned sequence of 500 bp results in 3 reads). The procedure was repeated for read lengths of 150, 300, 600, 1200 bp. An in previous studies, the simulated reads are contiguous substrings of the pruned sequences obtained without introducing sequencing errors (Matsen et al., 2010; Berger et al., 2011). This was motivated by the fact that errors may be treated in a pre-processing step, and that their frequency will tend to be reduced with the emergence of more accurate sequencing technologies (Salk et al., 2018; Matsen et al., 2010). A detailed description of the procedure can be found in Suppl. File S1.

### 2.5 Runtime and memory experiments

CPU times were measured for each software with the Unix command time, using a single CPU core and on the same desktop PC (64GB of RAM, Intel i7 3.6GHz, SSD storage). The time to execute the whole placement pipeline was measured. For EPA-RAxML, EPA-ng and PPlacer (the available likelihood-based software; see next section) this includes the compulsory alignment of QSs to the reference alignment. In the case of RAPPAS this includes the overhead induced by the initial loading of the pkDB into memory. The fast aligner program hmmalign from the HMMER package (Eddy, 2011) was used to align the QSs to the refA, as in previous placement studies (Matsen et al., 2010). Placement runtimes were measured for two of the reference datasets that we describe in the next section, chosen so as to represent different combinations of the two parameters with the largest impact on runtimes (number of taxa and alignment length). Each software was launched independently 3 times and the median CPU time was retained. For EPA-RaxML, EPA-ng and PPlacer, model parameters were manually defined to avoid any optimization phase prior to the placement. When available, masking options were kept as defaults. For EPA-RAxML and EPA-ng, a single thread was explicitly assigned (option -T 1). PPlacer only uses a single thread and required no intervention. Defaults values were used for all other parameters. RAPPAS is based on Java 8+, which uses a minimum of 2 threads. We ensured fair comparison by using the Unix controller cpulimit, which aligns CPU consumption to the equivalent of a single thread (options `--pid process_id --limit 100`). For each RAPPAS process, memory was monitored by programmatically reading its usage at regular intervals.

### 2.6 Methods and datasets used in experiments

The performance of RAPPAS (v1.00), both in terms of accuracy and computational efficiency, was compared against that of EPA (from RAxML v8.2.11) (Berger and Stamatakis, 2011), PPlacer (v1.1alpha19) (Matsen et al., 2010) and EPA-ng (v0.2.0-beta, with the agreement of its authors) (Barbera et al., 2018).

Reference datasets (refA, refT) used for evaluating accuracy were taken from (Matsen et al., 2010; Berger et al., 2011) with the exception of D155, which was built for this study and corresponds to a whole genome alignment and tree for the hepatitis C virus. Each dataset is identified by the number of taxa present in the tree (*e.g.* D150 has 150 taxa). In total, 10 reference datasets were tested. Descriptive metadata reporting the nature of the locus, the length of the alignment and the gap content are displayed in Figure 3 (left headers) and Suppl. File S2. One of the reference datasets (D140) is based on a reference alignment of protein sequences. Globally, the reference datasets represent a panel of loci commonly used in metabarcoding and metagenomic studies.

**Figure 3:**
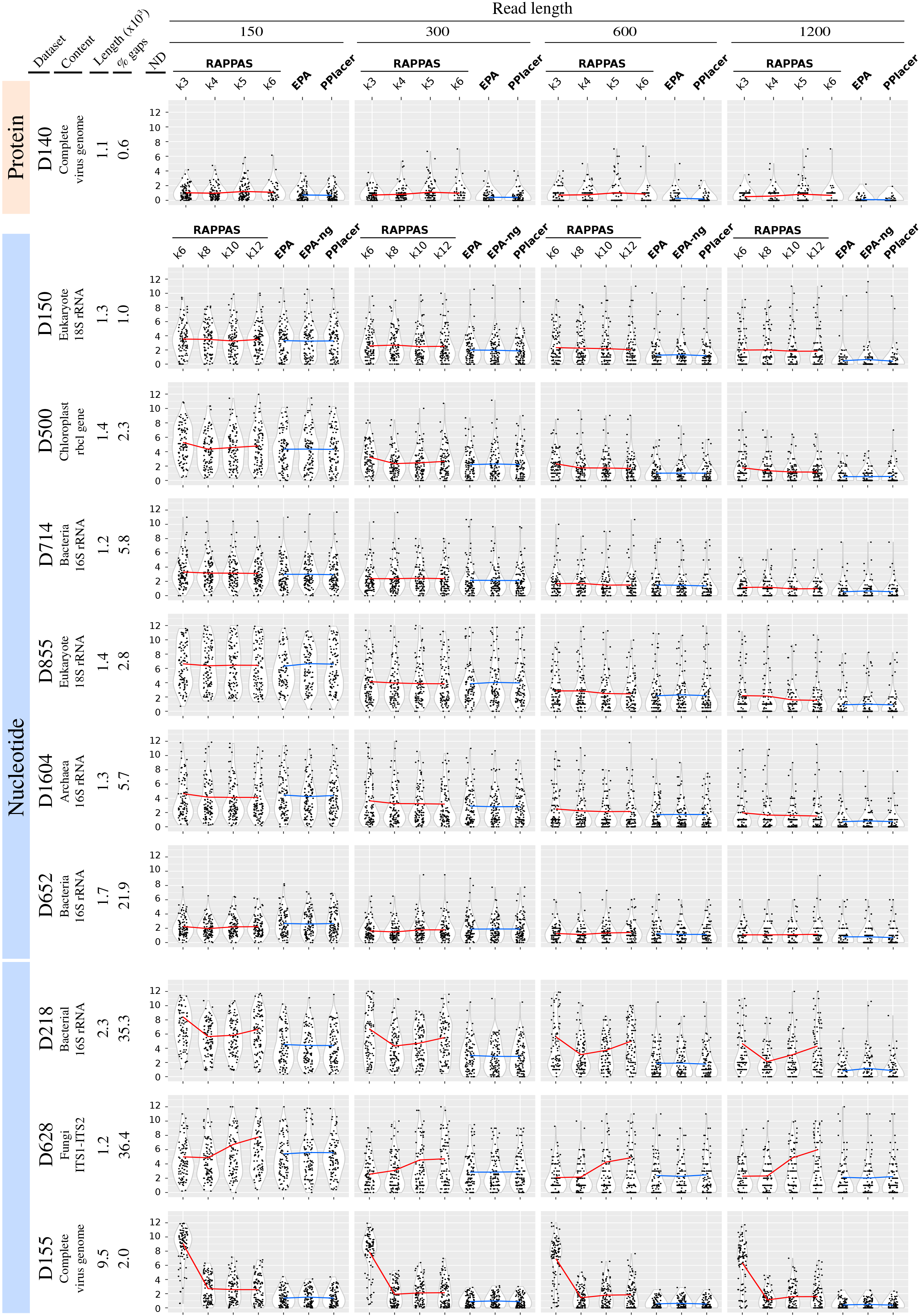
Accuracy comparison of RAPPAS and likelihood-based methods. Datasets corresponding to different marker genes or small genomes (left headers) were used to test placement accuracy for QSs of different lengths (top header). Lengths are measured in amino acids for D140 and in base pairs otherwise. Each grey box corresponds to a combination of dataset and QS length. In each box, the horizontal axis represents the tested software (RAPPAS was tested for several k-mer sizes), while the vertical axis corresponds to the measured node distance (ND) (the closer to 0, the more accurate the placement). For each software, black dots represent the mean NDs measured for 100 independent pruning experiments (see *Methods*) and white violins show their approximate distribution. Distribution means are linked by a red line (RAPPAS at different values of *k*) or a blue line (other software).

The runtime experiments are based on two of the above reference datasets, D652 (1.7 kbp) and D155 (9.5 kbp), and the QSs for placement were obtained in the following way. For D652, real-world 16S rRNA bacterial amplicons of about 150 bp were retrieved from the Earth Microbiome Project (Thompson et al., 2017) via the European Nucleotide Archive (Silvester et al., 2018) using custom scripts from github.com/biocore/emp/. Only sequences longer than 100 bp were retained. A collection of 3.1×10^8^ QSs were thus obtained. For D155, 10^7^ Illumina short reads of 150 bp were simulated from the 155 viral genomes in this reference dataset using the Mason read simulator (Holtgrewe, 2010). A SNP rate of 0.1% and very low indel rate of 0.0001% were chosen to avoid impacting negatively the runtime of the alignment phase for EPA, EPA-ng and PPlacer.

## 3 Results

### 3.1 Accuracy

The simulation procedure (see *Methods*) was repeated for 10 different reference datasets representing genes commonly used for taxonomic identification. Accuracy is measured on the basis of the *node distance* between observed placement and expected placement (*i.e.* the number of nodes separating the two edges). The closer to zero the node distances are, the more accurate the method is. Figure 3 summarizes the results of the experiments for each choice of dataset, read length, and software. Each point in the plot shows the mean node distance for the reads generated by one pruned subtree of the reference tree. Each violin plot is based on as many points as there are pruned subtrees (*i.e.* 100). Horizontal lines link the per software node distance means, calculated as the mean of the 100 means above. Detailed mean values are reported in Suppl. File S3.

As expected, given that they are based on the same general likelihood-based approach, EPA, EPA-ng, and PPlacer show very similar performance for all datasets and read lengths (Figure 3, blue lines almost perfectly horizontal). As expected, the longer the reads are, the more accurate the placement is (Figure 3, left to right boxes). The worst performances were observed for the D628 and D855 datasets, which agrees with previous studies (Matsen et al., 2010; Berger et al., 2011).

RAPPAS was subjected to the same tests using four k-mer sizes. The corresponding results can be classified into two groups. The first group contains the datasets for which using different values of *k* has a limited impact (Figure 3, all except the last three). For these, RAPPAS shows globally an accuracy comparable to likelihood-based software. In simulations based on the shortest read lengths (Figure 3, leftmost boxes) similar or even slightly better accuracies (leftmost boxes of D1604, D652) are measured. On the other hand, the longest read lengths generally resulted in lower accuracy compared to other methods (rightmost boxes of D140, D150, D500, D855, D1604).

The second group contains datasets where different values of *k* resulted in variable accuracy (Figure 3, D218, D628, D155). Two out of these three datasets are also characterized by a high proportion of gaps in their reference alignments (Figure 3, left headers), which may be an adverse factor for the current version of RAPPAS (discussed below). Finally, D155, which corresponds to the longest reference alignment (9.5 kbp), is the only dataset where shorter k-mer sizes clearly lowered the accuracy of RAPPAS.

### 3.2 Runtime

Figure 4 summarises runtimes measured for the placement of 10^3^ to 10^7^ QSs on two reference datasets corresponding to short and long reference sequences (a marker gene for bacteria, and a complete virus genome, respectively). To consider the actual time needed to place a set of metagenomic sequences, the runtime reported for EPA, EPA-ng, and PPlacer includes both read alignment and placement, while the runtime of RAPPAS includes the overhead related to loading the pkDB into memory.

**Figure 4:**
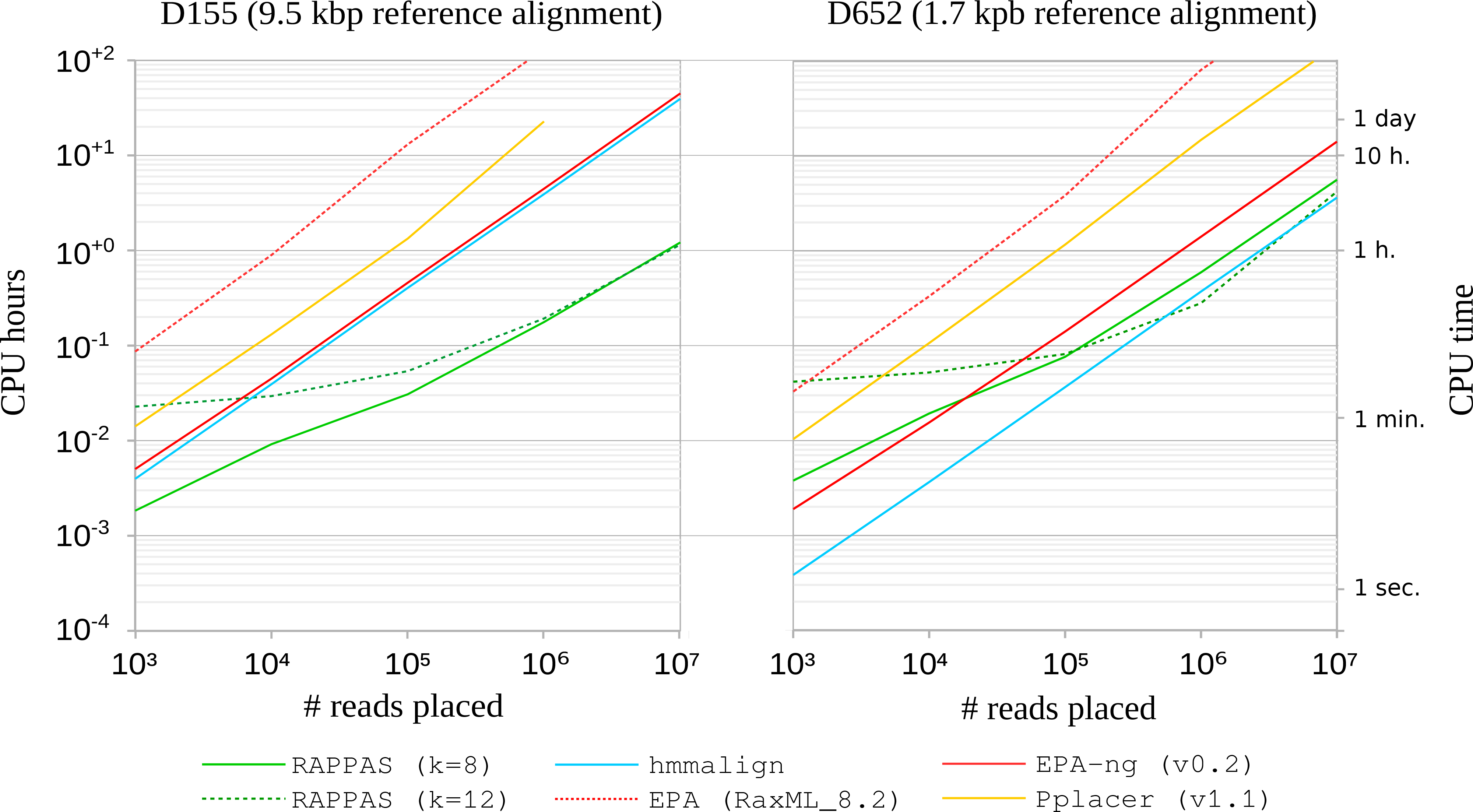
Placement runtime comparison. Runtimes were measured for a long (D155, left box) and a short (D652, right box) reference alignment, for a number of placement pipelines. The horizontal axis reports the number of placed reads. The blue line shows the time used to align those reads to the reference dataset, a preliminary step required by EPA, EPA-ng and PPlacer. For those pipelines, runtimes include the alignment phase (red and orange lines).

As expected, EPA, PPlacer, and EPA-ng show a runtime scaling linearly with the number of placed reads for both datasets. With the D155 dataset, EPA and PPlacer could not be run with a set of 10^7^ reads due to high memory requirements. Note that the runtime of EPA-ng for D155 is dominated by the time required to align the QSs (Figure 4, blue line in the left box).

The runtimes of RAPPAS appear to grow non-linearly when less than 10^5^ reads are placed due to the overhead imposed by loading the pkDB into memory. On the other hand, for larger read sets (≥10^5^) this overhead becomes negligible and the runtime of RAPPAS scales linearly with the number of reads. RAPPAS is much faster than the other methods. With D652, which is a short RefA (Figure 4, right box), RAPPAS was up to 5x faster than EPA-ng (5.0x for *k* = 12 and 2.4x for *k* = 8) and up to 53x faster than PPlacer (52.7x for *k* = 12 and 26.3x for *k* = 8). With D155, which is a longer refA (Figure 4, left box), greater speed increases are observed: RAPPAS was about 40x faster than EPA-ng (36.9x for *k* = 8 and 38.9x for *k* = 12) and about 130x times faster than PPlacer. Concretely, using a single CPU and a common desktop computer, RAPPAS is capable to place one million 100-150 bp DNA reads in 30 minutes on D652 and in 11 minutes on D155.

### 3.3 Implementation

RAPPAS is distributed as a standalone command-line application. It is compatible with any system supporting Java 1.8+. Ancestral sequence reconstruction is performed via an external program. Currently PhyML 3.3+ (Guindon et al., 2010) and PAML 4.9 (Yang, 2007) binaries are supported by RAPPAS. The main limiting factor for RAPPAS is the amount of memory required to build the pkDB (and subsequently, to load the pkDB for the placement phase). Figure 5 reports the memory and drive storage requirements related to building and storing the pkDB for three representative datasets and different values of *k*. D150 (few taxa, short alignment), D155 (few taxa, long alignment) and D1604 (many taxa, short alignment) show memory usage below 4GB for *k* = 10, and 12GB for *k* = 12, which indicates that RAPPAS can be launched on most standard laptop computers, if *k* is set to a low value. A Git repository, instructions to compile RAPPAS and tutorials are available at https://gite.lirmm.fr/linard/RAPPAS.

**Figure 5:**
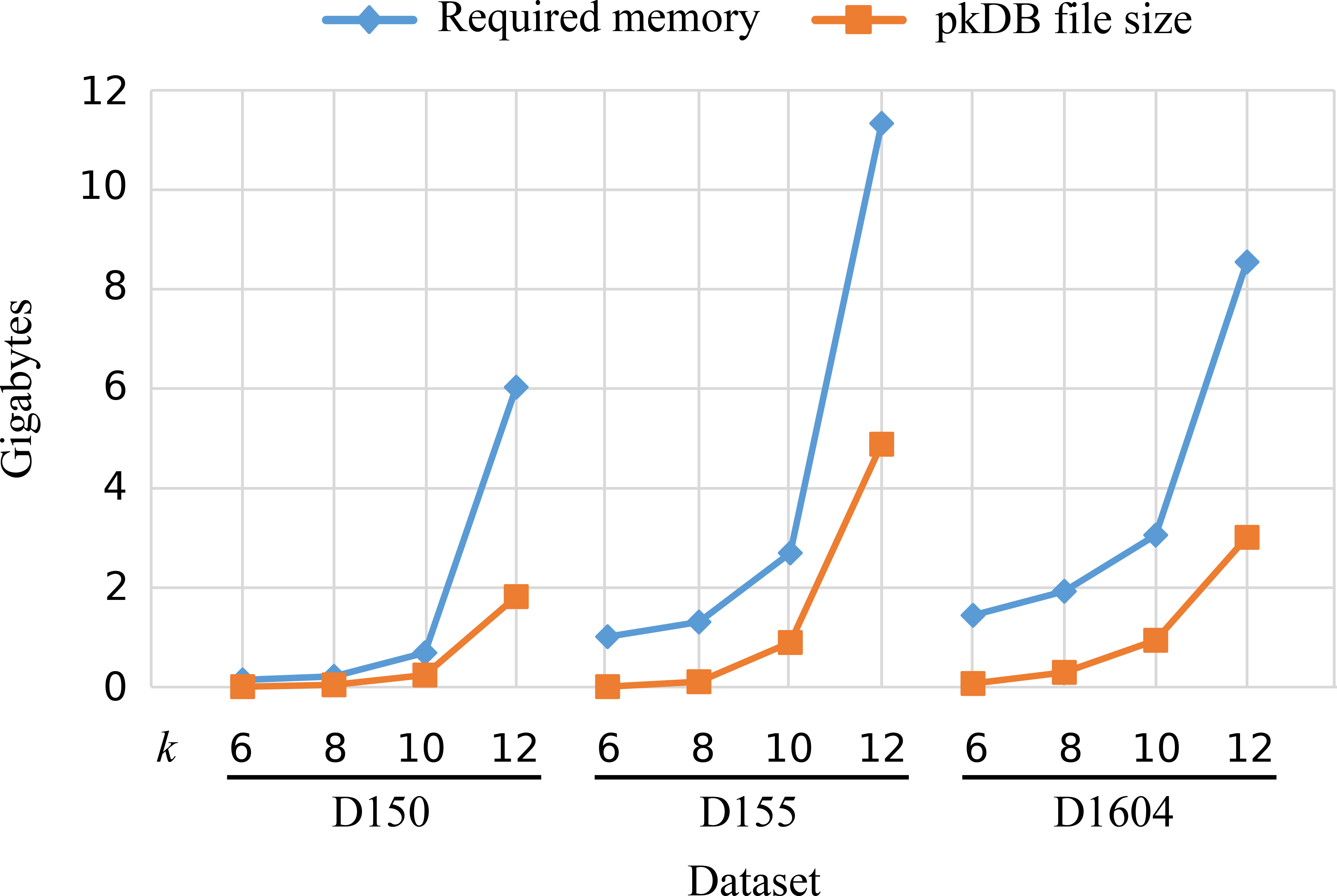
Memory and storage requirements related to the pkDB. Memory and storage requirements of RAPPAS are reported for three datasets, representative of different combinations of number of taxa and alignment length (D150, 1.3 kbp; D155, 9.5 kbp; D1604, 1.3 kbp). The blue line reports the memory peaks experienced during pkDB construction. The orange line displays the size of the pkDB when saved as a compressed binary file.

## 4. Discussion

### 4.1 Advantages of alignment-free placements

The novel algorithm introduced in RAPPAS removes a major limitation imposed by previous PP software; these methods rely on the computation of a multiple alignment merging the reference alignment and query sequences, a step which must be repeated for each new set of query sequences (Figure 2-A). The idea of attaching k-mers to pre-computed metadata to accelerate sequence classification has been explored by algorithms combining taxonomically-informed k-mers and LCA-based classification (Liu et al., 2018; Müller et al., 2017; Wood and Salzberg, 2014; Ounit et al., 2015). However, none of these methods have the goal of producing evolutionarily precise classifications based on probabilistic models of sequence evolution. RAPPAS adapts this idea to PP by reconstructing sets of plausible, phylogenetically-informed k-mers, called phylo-kmers.

Compared to previous PP methods, RAPPAS has two critical algorithmic advantages (see Suppl. File S1 for details). First, its runtime does not depend on the length of the reference alignment, since the sequences in the reference tree, and those that are related to them, are summarized by their probable k-mer content. Note that the alignment phase required by the other PP methods has a runtime that scales linearly with the length of refA (if profile alignment is used (Eddy, 1998)). Second, while the placement runtime of other PP methods is linear in the number of reference taxa (*i.e.* the size of refT) (Matsen et al., 2010), for RAPPAS it depends only on the number of edges associated to the queried phylo-kmers, which may be significantly smaller than the number of taxa. This second property has limited impact for a small refT with very similar reference sequences, or for small values of *k* (as k-mers will be assigned to many tree edges), but is favorable for a large refT, spanning a wide taxonomic range, and large values of *k* (as k-mers will be assigned to a low proportion of the edges).

Our experiments on two reference datasets with very different combinations of the parameters governing runtimes (number of taxa and alignment length) show that, on real data, the initial implementation of RAPPAS is already faster than alternative PP pipelines (Figure 4). This includes EPA-ng that, despite being based on low level optimizations (e.g. CPU instructions, optimized libraries, etc.) (Barbera et al., 2018), is only faster than RAPPAS when few query sequences are analysed. As expected, the speed gain provided by the alignment-free approach appears to be more pronounced for long reference alignments (Figure 4, left box). Finally, while the present implementation is based on a single thread, most steps of RAPPAS should be fully parallelizable, offering even faster placements in the future.

### 4.2 Accuracy limitations

The presence of dense gappy regions in refA appears to have a negative impact on the accuracy of RAPPAS, especially in combination with larger k-mer sizes: D218 and D628 are the most gappy of the tested datasets and cause increasingly worse accuracy (compared to EPA/PPlacer) when *k* increases from 8 to 12 (Figure 3). The likely reason behind this limitation is the following. As is standard practice in phylogenetics, substitution models used during ancestral state reconstruction do not treat gaps in refA as sequence states. As a result, our phylo-kmers are sequences composed of nucleotides (or amino acids) only and cannot be used to model the presence of indels in the query sequences. The by-default construction of phylo-kmers described in the Methods produces phylo-kmers that will not match the query sequences having indels among the corresponding positions. Clearly, the longer the k-mers are, the more likely these failed matches are to occur. A possible solution to this issue consists of recording the coordinates of the indels present in refA, and generate phylo-kmers that skip the refA intervals defined by those coordinates. This solution is implemented in RAPPAS, and we describe it in more detail in Suppl. File S1. Other improvements may be possible, for example allowing inexact (approximate) matching between the query and the stored k-mers, as explored in several k-mer based sequence classification tools (Müller et al., 2017; Wood and Salzberg, 2014) or in tools for sequence comparison (Horwege et al., 2014). These or other solutions, adapted to a probabilistic setting, will be examined in the future to reduce the sensitivity of RAPPAS to gaps.

Another limitation of RAPPAS is that although it is practically as accurate as likelihood-based methods for read lengths that are commonly produced by Illumina sequencing (*e.g.* 150-300 bp), it tends to become less accurate on longer reads (Figure 3). This is not surprising: query sequences are treated by RAPPAS as unordered collections of k-mers, meaning that RAPPAS does not evaluate whether the matches between the phylo-kmers and the query k-mers respect some form of positional consistency or collinearity. It is intuitive that this problem is potentially more harmful to accuracy for longer query sequences. Since long read sequencing is developing rapidly, future versions of the algorithm will deal with this issue by taking into account the positions within refA from which the phylo-kmers originated.

We note that low values of *k* and long reference alignments make it possible for the same phylo-kmer to be generated for a single edge at more than one position in refA. In this case, only the highest probability among these positions is stored in the pkDB (see Methods). Frequent occurrence of this event may have an undesired impact on the placement score of reads, and ultimately affect placement accuracy. This is the likely reason for the lower accuracy observed in the D155 dataset (especially for *k* = 6), which corresponds to a complete viral genome rather than a single short genetic marker (Figure 3, lowest row).

### 4.3 Applying RAPPAS to “portable” metagenomics

Improvements impacting the computational efficiency (and not just accuracy) of RAPPAS are also conceivable. For example, tailored k-mer indexing techniques can be developed similarly to other recent work (Müller et al., 2017; Liu et al., 2018), and the memory footprint of the pkDB can be reduced by limiting storage to the most discriminant phylo-kmers (Ounit et al., 2015). Despite the many potential improvements to RAPPAS, it is already faster than other PP implementations on real datasets (Figure 4).

The design principle behind RAPPAS (Figure 1) is particularly adapted to metagenomic analyses run on portable devices with reduced computing capacity. As a preparatory step, one could build a set of pkDBs from reference datasets maintained in standard genetic marker databases such as EukRef (eukref.org), SILVA (Quast et al., 2013), RDP (Cole et al., 2014), PhytoRef (Decelle et al., 2015), or more specialized reference datasets maintained locally. These pkDBs can then be shared on portable computers equipped with portable sequencing devices. RAPPAS would place many reads on the different reference trees, efficiently identifying a sample’s composition in the phylogenetic groups of interest. This approach has a strong potential in the context of field-based DNA sequencing, which is rapidly developing thanks to the increasing portability of sequencing technologies (Lu et al., 2016; Edwards et al., 2016; Batovska et al., 2017). As the RAPPAS algorithm evolves to handle longer query sequences with increased accuracy and efficiency, it will open the door to phylogenetic placement as a means to real-time onsite species identification.

## Acknowledgements

We would like to thank Olivier Gascuel for helpful advice, as well as Anne-Mieke Vandamme, Krystof Theys, Pieter Libin, Frédéric Mahé and Jakub Truszkowski for sharing data and fruitful discussions. BL is also grateful to Vincent Lefort for his everyday advice. BL, KS and FP conceived of the study; BL, KS and FP designed the algorithm; BL coded the software; BL, KS and FP wrote the manuscript.

